# Unveiling the Role of Endoplasmic Reticulum Stress Pathways in Canine Demodicosis

**DOI:** 10.1101/2023.08.11.552979

**Authors:** Pamela A Kelly, Gillian P. McHugo, Caitriona Scaife, Susan Peters, M. Lynn Stevenson, Jennifer S McKay, David E. MacHugh, Irene Lara Saez, Rory Breathnach

## Abstract

Canine demodicosis is a prevalent skin disease caused by overpopulation of a commensal species of *Demodex* mite, yet its precise cause remains unknown. Research suggests that T cell exhaustion, increased immunosuppressive cytokines, induction of regulatory T cells, and increased expression of immune checkpoint inhibitors may contribute to its pathogenesis. This study aimed to gain a deeper understanding of the molecular changes occurring in canine demodicosis using mass spectrometry and pathway enrichment analysis. The results indicate that endoplasmic reticulum stress is promoting canine demodicosis through regulation of three linked signalling pathways: eIF2, mTOR, and eIF4 and p70S6K. These pathways are involved in the modulation of Toll-like receptors, most notably TLR2, and have been shown to play a role in the pathogenesis of skin diseases in both dogs and humans. Moreover, these pathways are also implicated in the promotion of immunosuppressive M2 phenotype macrophages. Immunohistochemical analysis, utilizing common markers of dendritic cells and macrophages, verified the presence of M2 macrophages in canine demodicosis. The proteomic analysis also identified immunological disease, organismal injury and abnormalities, and inflammatory response as the most significant underlying diseases and disorders associated with canine demodicosis. This study demonstrates that *Demodex* mites, through ER stress, unfolded protein response and M2 macrophages contribute to an immunosuppressive microenvironment thereby assisting in their proliferation.

## Introduction

Canine demodicosis is a prevalent skin disease commonly seen in primary care small animal practice(1, 2). It occurs when a species of the commensal skin mite *Demodex* genus overpopulates the skin (3). There are three identified species of *Demodex* mites reported in dogs, with the most common being *Demodex canis*(2, 4). The other species include *Demodex injai* and *Demodex cornei*, the latter is a likely variant of *Demodex canis*(5–8). The disease is clinically classified based on lesion location and extent (generalized, localized) and or age of onset (juvenile onset, less than 18 months of age, and adult onset, greater than four years of age)(1, 2). The adult-onset disease is typically linked with concurrent illnesses or treatments that cause immunosuppression, such as neoplasia, endocrinopathies, and corticosteroid administration (2, 3, 9, 10).

The exact pathogenesis of canine demodicosis is not yet fully understood. The leading hypothesis is of immune dysregulation and T cell exhaustion, with dogs with demodicosis having increased expression of immunosuppressive cytokine IL-10, decreased expression of IL-2, IL-21 and reduced circulating CD4+ cells. This hypothesis is supported by the high incidence of demodicosis in dogs with immunosuppressive diseases or undergoing immune modulating therapy(2, 4, 10–12). However, there are instances where no evidence of immunosuppression exists, and in these cases, appropriate antiparasitic treatment leads to recovery and limited relapse (2, 9, 13–15). With these cases it is hypothesised that the *Demodex* mite itself is modulating the immune response. Research has shown that TLR2 expression is high in cases of canine demodicosis, and it is thought to be one of the main contributing causes of the high levels of IL-10. Recently it has been demonstrated that Regulatory T cells, known to produce IL-10, are increased in the skin of dogs with demodicosis in comparison with healthy canine control skin. Increased gene expression of several immune check point molecules including PD-1/PD-L1 and CTLA-4 has also recently been shown in cases of canine demodicosis^49^. These findings support the possibility that the *Demodex* mites could induce changes in the hair follicle, sebaceous gland, and or immune cells that promote immune tolerance and support its proliferation.

Research into the pathogenesis of canine demodicosis has mainly focused on mRNA expression changes in both blood and tissue, with only a few studies exploring changes at the protein level using immunohistochemistry and flow cytometry(13, 14, 16–26). Protein-level studies have primarily focused on assessing a limited panel of proteins such as cytokines and cell receptors(16–18, 20, 22). Proteomics is a crucial technology in human medicine for understanding disease pathogenesis, discovering biomarkers, and identifying potential treatment targets(27). Proteomics offers advantages over conventional genomics and transcriptomics tools as it can detect post-translational modifications like phosphorylation, glycosylation, and acetylation, providing a deeper characterization of skin diseases’ pathogenesis (28). Currently, there are limited proteomic profiling studies reported in veterinary medicine for skin diseases.

The aim of this study was to enhance our understanding of the molecular changes occurring in canine demodicosis. To achieve this, we examined the proteomic profile of formalin-fixed paraffin-embedded skin samples using mass spectrometry (MS) of dogs with canine demodicosis and compare with profiles from healthy control skin. Additionally, we aimed to characterise the dendritic cell and macrophage population within lesional skin of canine demodicosis using immunohistochemical analysis.

## Methods

### Proteomics

#### Case selection

This study obtained ethical exemption by the Animal Research Ethics Committee (AREC) at University College Dublin (AREC E 19 09 Kelly).

Ten formalin fixed paraffin embedded (FFPE) skin samples from dogs diagnosed histologically with canine demodicosis were selected. All ten had skin swabs for microbiological assessment taken on the day of biopsy sampling of which culture results were considered ‘normal skin flora’ (29). Details of signalment and microbiological culture are available in the supplementary material (Supplementary Table 1). Ten FFPE skin samples from dogs that had no history, gross or histological evidence of skin disease were selected as controls. Details of signalment of the control cases is available in supplementary material (Supplementary Table 1).

#### Protein extraction

Up to three scrolls from each FFPE block, with a thickness of up to 15 μm and a tissue area of up to 100 mm^2^ were trimmed using a microtome and placed into sterile 2 ml collection tubes. Samples once cut were kept at -80°C until they could be deparaffinised. Deparaffinisation was carried out using heptane and methanol. Protein exaction from the deparaffinised sample was carried out using the Qiagen Qproteome kit (Qiagen, Hilden, Germany) as per the manufacturer’s protocol. Briefly, following deparaffinisation, extraction buffer (EXB plus, Qiagen) supplemented with β-mercaptoethanol was added and the samples were heat treated. In preparation for MS the protein was isolated using chloroform, methanol, and ddH_2_O separation. The protein samples were then dissolved in 1% (w/v) RapiGest (Waters Corp, Etten-Leur, Netherlands) and were digested to peptides using a trypsin digest as indicated in the Qproteome kit protocol.

#### Mass Spectrometry

Following trypsin digestion, the peptides were cleaned using C18 ZipTip® (Merck Millipore, Massachusetts, United States). Samples were run on a Bruker timsTOF Pro mass spectrometer (Bruker Daltonics, Bremen, Germany) connected to an Evosep One system (EvoSep BioSystems, Odense, Denmark). Tryptic peptides were loaded onto Evotips and separated on a reversed-phase C18 Endurance column (15 cm × 150 μm ID, C18, 1.9 μm) using the pre-set 30 SPD method. Mobile phases were 0.1% (v/v) formic acid in ddH_2_O (phase A) and 0.1% (v/v) formic acid in acetonitrile (phase B). The peptides were separated by an increasing gradient of mobile phase B for 44 min using a flow rate of 0.5 µL/min.

The mass spectrometer was operated in positive ion mode with TIMS (Trapped Ion Mobility Spectrometry) and PASEF (Parallel Accumulation Serial Fragmentation) enabled. The accumulation and ramp times for the TIMS were both set to 100 ms, with an ion mobility (1/k0) range from 0.6 to 1.6 Vs/cm. A scan range of (100-1700 m/z) was performed at a rate of 10 PASEF MS/MS frames to 1 MS scan with a cycle time of 1.17s.

#### Proteomic Data Analysis

Protein identification and label-free quantification (LFQ) normalization of MS/MS data was performed using MaxQuant v2.0.3.0 (www.maxquant.org)(30). Variable modifications selected were Acetyl (Protein N-term) and Oxidation (M), while trypsin was selected as the digestion enzyme and the maximum number of allowed missed cleavage was two. MS/MS data were correlated against the *Canis lupus familaris* reference proteome downloaded from Uniprot (January 2023) using the Andromeda search algorithm incorporated in MaxQuant software, including a contaminant sequence set. Data analysis, processing and visualization were performed using Perseus v.2.1.4.0 (www.maxquant.org) following standard steps for LFQ analysis(31). Briefly, LFQ normalized peptide intensity values from the MaxQuant analysis were used to quantify protein abundance. Data were filtered to remove protein groups that were only identified by peptides that carry one or more modified amino acids, those matching to the reverse database, and those identified as potential contaminants. Then log_2_ transformation was performed, with subsequent grouping of samples according to aetiology (control, demodicosis). Further filtering was performed whereby only proteins present in at least 70% of samples were retained, and a two-sample t-test was performed. Statistical testing was done at the two-tailed α level of 0.05 (*P* < 0.05) to identify significantly differentially abundant proteins across the two aetiologies. In addition, we also used the permutation-based *q* false discovery rate (FDR) method to adjust *P* values and correct for multiple testing (32, 33). A *q* value threshold of 0.01 was used to further filter differentially abundant proteins.

#### Metascape® analysis

Metascape® is an online software based on the OMICS database(34). It has functions such as functional enrichment, interactome analysis, gene annotation, and membership search, and it can analyse and annotate given genes. The Metascape® (v3.5.20230501) express analysis function was used to analyse the functions and pathways of the protein groups identified as either in significantly high abundance or lower abundance. The *q* false discovery rate (FDR, 0.01) method was applied to filter the input gene symbols. Only protein groups in which gene symbol IDs were available were used for the gene ontology (GO) overrepresentation analyses. The set of background genes used were the gene IDs of proteins found across both the demodicosis and control groups.

#### Ingenuity® Pathway Analysis

Ingenuity® Pathway Analysis (IPA) software (35) was used with the Ingenuity® Knowledge Base (Qiagen, Redwood City, CA, USA; release date December 2022) to identify enriched canonical pathways for the differentially abundant proteins that had corresponding gene symbols. IPA® Core Analysis was performed using the default settings with the user data set as the background, high predicted confidence and all nodes selected. As with the Metascape® analysis, *q* false discovery rate (FDR, 0.01) method was applied to filter the input gene symbols. For identification of overrepresented canonical pathways, a stringent Benjamini–Hochberg (B-H) *P*-value adjustment was also applied with a B-H FDR *P*_adj_ threshold < 0.05(36).

### Immunohistochemistry

#### Case selection

Ten FFPE skin samples from dogs diagnosed histologically with canine demodicosis were selected. A further ten FFPE skin samples from dogs that had no history, gross or histological evidence of skin disease were selected as controls. Details of signalment of cases is available in supplementary material (Supplementary Table 2).

#### Immunohistochemistry

The antibodies applied are common cell markers used to identify different lineages of macrophage and dendritic cells in tissues. Detailed methods for immunohistochemistry are available in the supplementary material (Supplementary Material 1). Briefly, 5-μm-thick sections were prepared from the FFPE tissue blocks for each case, rehydrated, and stained by Immunoperoxidase methods as indicated in Table 1. 3,3′-Diaminobenzidine (DAB) was used as the chromogen and Gill’s haematoxylin as counterstain.

**Table 1:** Antibodies and immunohistochemical procedures.

| Antibody | Dilution | Incubation | Pretreatment | Source |
| --- | --- | --- | --- | --- |
| IBA-1 | 1:1500 | 30 minutes | HIER, citrate buffer, pH 6 | Wako Chemical |
| CD301 | 1:50 | 30 minutes | HIER, citrate buffer, pH 6 | Invitrogen |
| CD90 | 1:100 | 60 minutes | Proteinase K | Novus Biologicals |
| CD163 | 1:250 | 30 minutes | DIVA | Novus Biologicals |
| E-Cadherin | 1:200 | 30 minutes | DIVA | Cell Signaling Technology |
| CD204 | 1:100 | 30 minutes | Proteinase K | Novus Biologicals |

#### Evaluation of immunostaining

Using an Olympus BX41 microscope (Olympus Optical Co., Tokyo, Japan) connected to a digital camera (SC50, Olympus), five representative areas were selected that showed the highest population of positive immunolabelled cells within each tissue section. These were imaged using the 400× power objective and saved as JPEG files. To allow for comparison of immunolabelling across multiple skin samples, image acquisition was performed in parallel for the entire set, using identical settings and exposure times. The image analysis software ImageJ (version 1.54d) was used to quantify the expression of each marker using the IHC profiler plugin(37, 38). This IHC analysis tool uses colour deconvolution and computerized pixel profiling leading to the assignment of automated scores to the respective image of % high positive, % positive, % low positive and % negative stained cells. The high positive, positive, and low positive scores were combined to provide a total percentage score of immunolabelled positive cells within an image. This percentage total positive scores were used for statistical analysis. The Student’s t-test was use to compare the means between the demodicosis group and the control group (*P* < 0.05).

## Results

### Proteome profile

The general proteome profiling of the analysed skin samples was examined to identify differences in protein group paucity or abundance. We identified 1,123 protein groups across all samples (Supplementary Table 3), after removing proteins identified by site, matching to the reverse database and contaminants. Following further filtering for at least 70% of valid values (i.e., removing protein groups with <70% valid values) a total of 942 protein groups were found across the samples. Principal Component Analysis (PCA) was performed based on the protein abundance values from LC-MS/MS. A plot of the first two principal components (PC1 and PC2) (Figure 1) showed clear differentiation and clustering of samples within the canine demodicosis (*n* = 10) and control (*n* = 10) groups, and with 24.5% and 12.5% of the variation explained by PC1 and PC2, respectively. Filtering by the FDR-adjusted *P*-value (*q* < 0.01), showed that 267 protein groups were significantly differentially expressed between the two groups (Figure 2; Supplementary Table 4). Of these protein groups, 154 were observed to be more highly and 113 more lowly abundant, respectively in the demodicosis group compared to the control group. The 20 most abundant and 20 least abundant proteins for this contrast are detailed in Tables 2 and 3.

**Figure 1:**
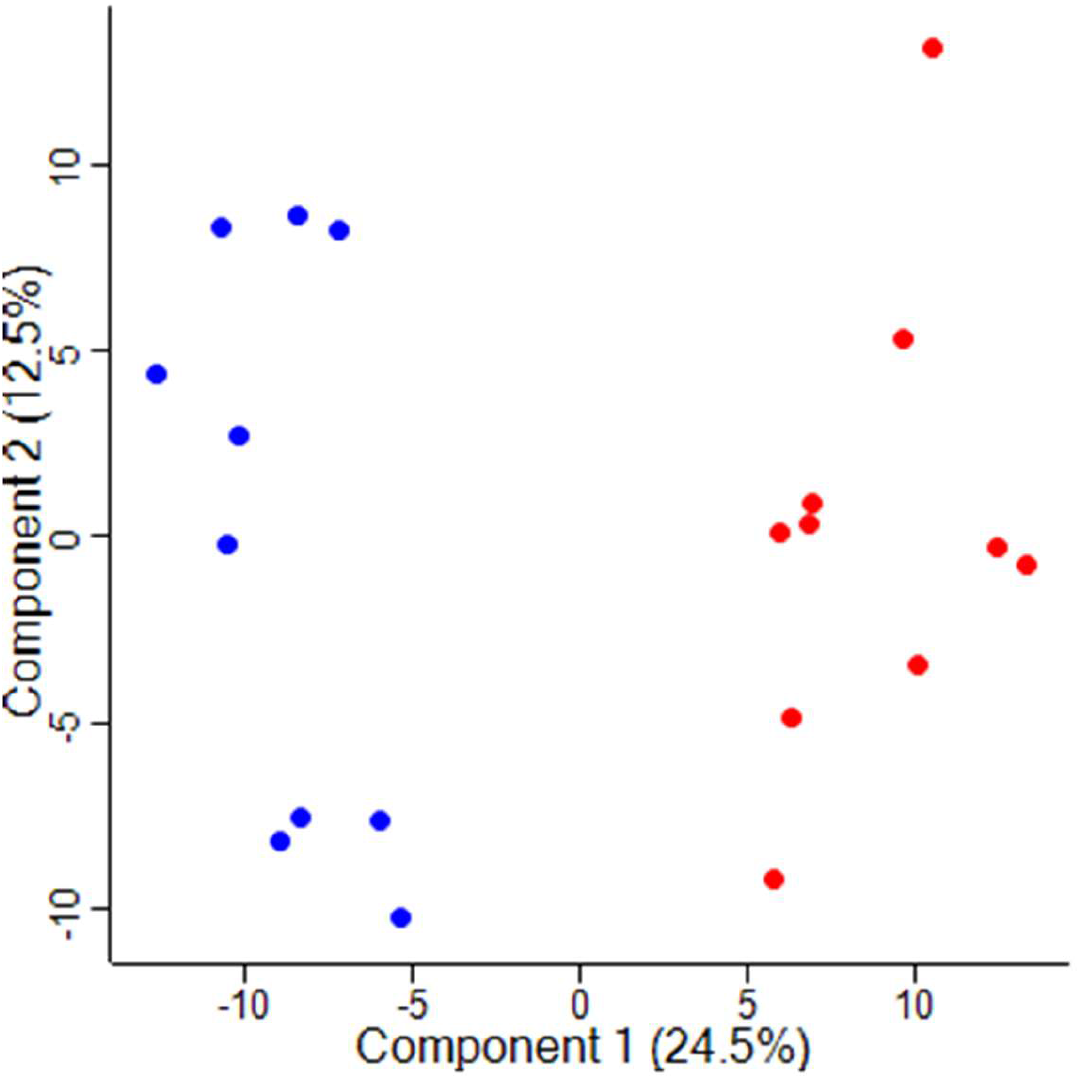
Principal component analysis (PCA). Each data point represents a sample with control samples shown in blue and demodicosis samples shown in red.

**Figure 2:**
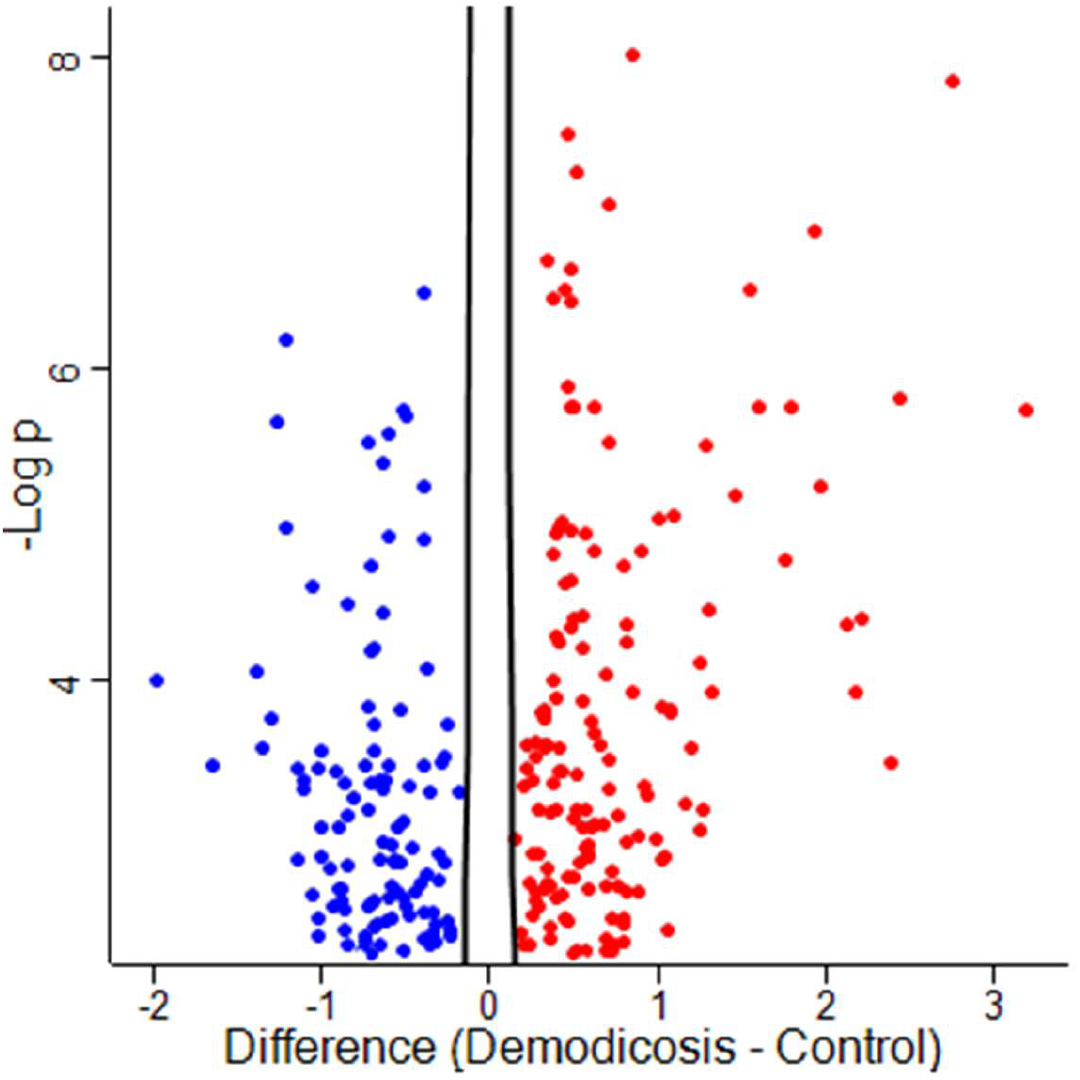
Volcano plot of proteins found to be significantly differentially abundant in the demodicosis group compared with the control group (*q* < 0.01). Proteins with significantly decreased abundance are shown in blue and proteins with significantly increased abundance are shown in red.

**Table 2:** Table of the twenty most statistically significant proteins displaying increased abundance the demodicosis case group compared with the control group (*q* < 0.01).

| Protein name | Gene | Location | q(adjusted *P*)-value |
| --- | --- | --- | --- |
| Myeloperoxidase | MPO | Cytoplasm | 0.00001 |
| Protein S100A12 | S100A12 | Cytoplasm | 0.00001 |
| Cornifin-like | LOC119868937 | Cytoplasm | 0.00001 |
| EF-hand domain-containing protein | S100A9 | Cytoplasm | 0.00001 |
| Coronin | CORO1A | Cytoplasm | 0.00001 |
| Major histocompatibility complex, class II, DQ alpha 1 | DLA-DQA1 | Plasma Membrane | 0.00001 |
| Cathepsin C | CTSC | Cytoplasm | 0.00001 |
| Cathepsin S | CTSS | Cytoplasm | 0.00001 |
| Glia maturation factor | GMFG | Cytoplasm | 0.00001 |
| Ras homolog family member | RHOG | Cytoplasm | 0.00001 |
| Lymphocyte cytosolic protein 1 | LCP1 | Cytoplasm | 0.00001 |
| Profilin | PFN1 | Cytoplasm | 0.00001 |
| PYD and CARD domain containing | PYCARD | Cytoplasm | 0.00001 |
| Actin-related protein 3 O | ACTR3 | Plasma Membrane | 0.00001 |
| Actin-related protein 2/3 complex subunit 4 | ARPC4 | Cytoplasm | 0.00001 |
| 60S acidic ribosomal protein P2 | RPLP2 | Cytoplasm | 0.00001 |
| 40S ribosomal protein SA | RPSA | Cytoplasm | 0.00001 |
| B-cell antigen receptor complex-associated protein alpha chain | CD79A | Plasma Membrane | 0.00001 |
| 60S ribosomal protein L9 | RPL9 | Nucleus | 0.00001 |
| Enolase 1 | ENO1 | Cytoplasm | 0.00001 |
| Chloride intracellular channel protein | CLIC1 | Nucleus | 0.00001 |
| 60S acidic ribosomal protein P0 | LOC481399 | Cytoplasm | 0.00001 |
| 40S ribosomal protein S8 | RPS8 | Cytoplasm | 0.00001 |
| Phosphoglycerate mutase | PGAM1 | Cytoplasm | 0.00001 |
| Serine/arginine-rich splicing factor 1 | SRSF1 | Nucleus | 0.0001 |

**Table 3:** Table of the twenty most statistically significant proteins displaying decreased abundance the demodicosis case group compared with the control group (*q* < 0.01).

| Protein name | Gene | Location | q(adjusted P)-value |
| --- | --- | --- | --- |
| Acyl-CoA dehydrogenase very long chain O | ACADVL | Cytoplasm | 0.00001 |
| Actinin alpha 1 | ACTN1 | Cytoplasm | 0.00001 |
| Basal cell adhesion molecule | BCAM | Plasma Membrane | 0.0001 |
| Caveolae associated protein 1 | CAVIN1 | Nucleus | 0.00018 |
| Carboxylesterase 5A | CES5A | Extracellular Space | 0.00073 |
| EH domain containing 2 | EHD2 | Nucleus | 0.000082 |
| Fetuin B | FETUB | Extracellular Space | 0.00072 |
| Four and a half LIM domains 1 | FHL1 | Cytoplasm | 0.000451 |
| Filamin C | FLNC | Cytoplasm | 0.000438 |
| Guanine deaminase | GDA | Cytoplasm | 0.000585 |
| Glutathione S-transferase mu 3 | GSTM3 | Cytoplasm | 0.00001 |
| Isocitrate dehydrogenase [NADP] | IDH1 | Cytoplasm | 0.000091 |
| Keratin 4 | KRT77 | Cytoplasm | 0.000636 |
| IF rod domain-containing protein | KRT86 | Cytoplasm | 0.000108 |
| Laminin subunit beta 2 | LAMB2 | Extracellular Space | 0.000093 |
| Laminin subunit gamma 1 | LAMC1 | Extracellular Space | 0.00001 |
| ATP synthase subunit b | LOC119872230 | Mitochondria | 0.00001 |
| Malate dehydrogenase | MDH1 | Cytoplasm | 0.000343 |
| Myosin heavy chain 14 | MYH14 | Extracellular Space | 0.000176 |
| Nidogen 1 | NID1 | Extracellular Space | 0.00001 |
| Podocan | PODN | Cytoplasm | 0.0000076 |
| Pre-mRNA processing factor 8 | PRPF8 | Nucleus | 0.0006 |
| Transmembrane protein 43 | TMEM43 | Nucleus | 0.00001 |
| Tropomyosin 1 | TPM1 | Cytoplasm | 0.000143 |
| Vinculin | VCL | Plasma Membrane | 0.00001 |

### Pathway analysis

Results from the Metascape® gene ontology (GO) overrepresentation analysis of 131 highly and 110 lowly abundant proteins (with available gene symbol IDs) are shown in Figure 3. The analysis used the proteins observed across all samples (1123 proteins, with 1020 gene symbol IDs available) as the background set for the overrepresentation analysis.

**Figure 3:**
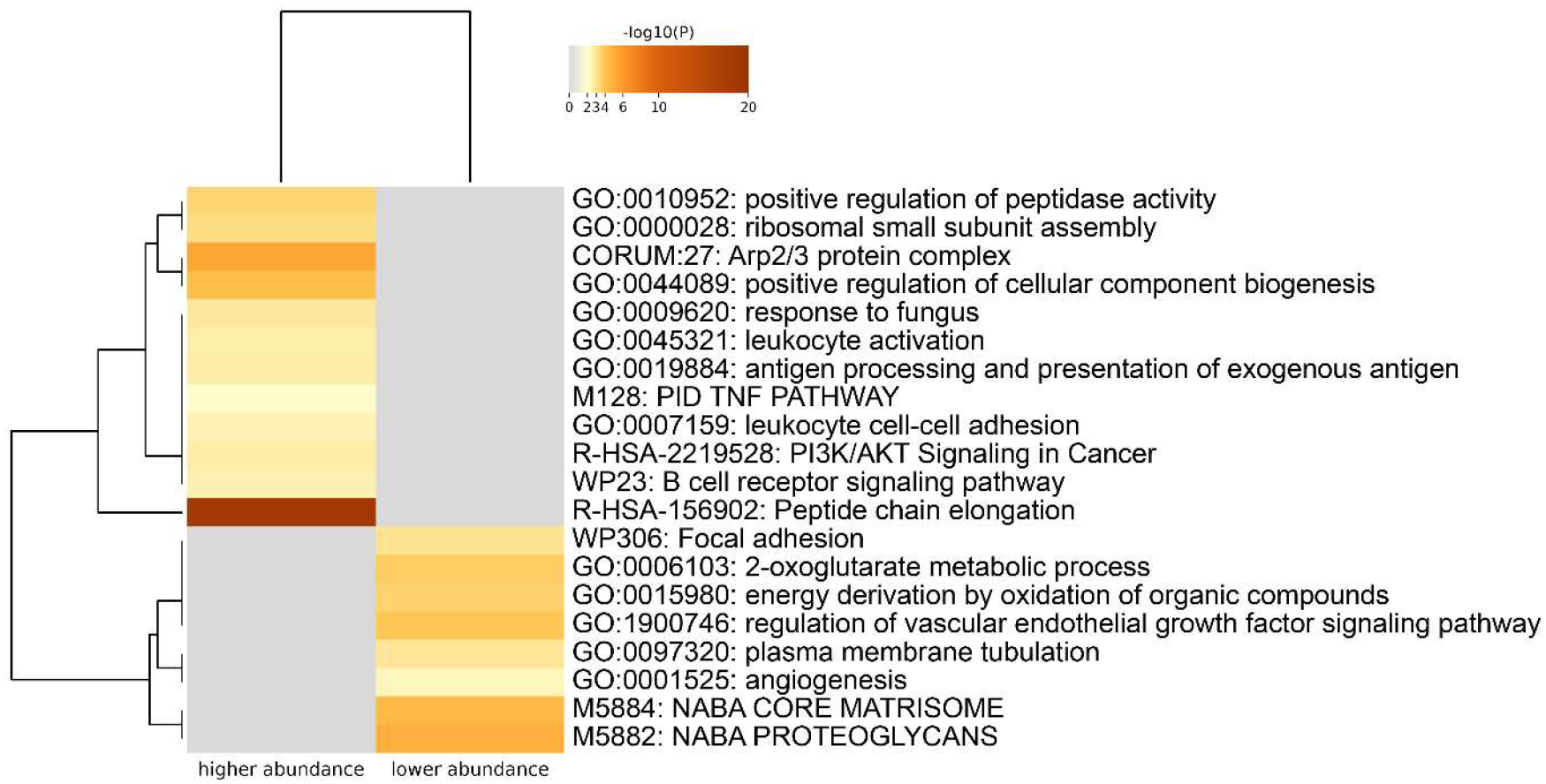
Bar chart showing the enrichment ontology clusters obtained by Metascape® enrichment analysis from the most significant high abundance and low abundance proteins found in the canine demodicosis group compared with the control group.

From the set of 1,123 detectable protein groups found across all samples, 1,080 had corresponding gene symbol IDs and 1015 of these could be mapped within IPA®. The majority of those unmapped (55/65) were LOC gene symbols for which a formal ID has not yet been determined. The B-H FDR *P*_adj._ (*q*) threshold < 0.01 gave 238 gene symbol IDs for the IPA analysis, 132 of which had an increased log_2_ fold change (log_2_FC) and 106 of which had a decreased log_2_FC against a background data set of 1014 analysis-ready gene symbols, 507 of which had an increased log_2_FC and 507 of which had a decreased log_2_FC. With a B-H FDR *P*_adj._ threshold < 0.05 there were four statistically significant enriched IPA® canonical pathways: *eIF2 Signaling, Coronavirus Pathogenesis Pathway, mTOR Signaling*, and *Regulation of eIF4 and p70S6K Signaling* (Figure 4). A schematic diagram of each of these pathways highlighting the differentially expressed molecules is provided in Supplementary Figures 1-4.

**Figure 4:**
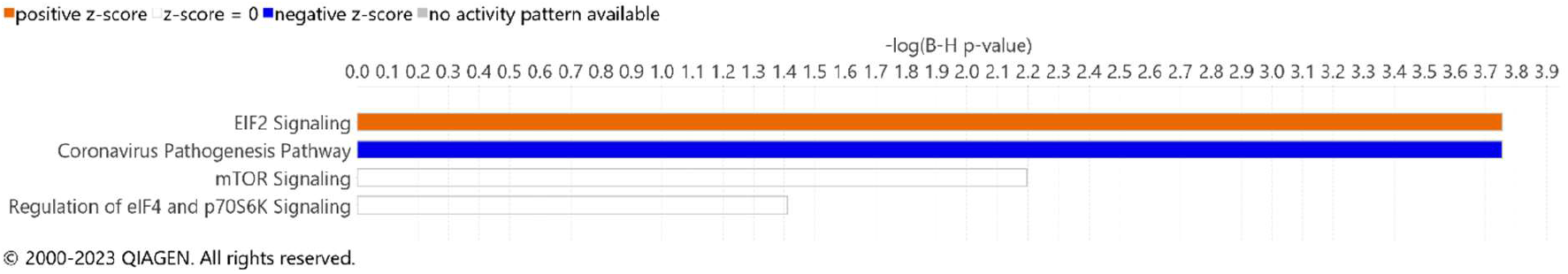
Bar chart showing the four statistically significant enriched IPA® canonical pathways in order of decreasing -log (B-H FDR-*P*_adj._): *EIF2 Signaling*, *Coronavirus Pathogenesis Pathway*, *mTOR Signaling*, and *Regulation of eIF4 and p70S6K Signaling*. The orange and blue colours of the bars indicate the predicted pathway activation or inhibition, respectively. White bars indicate pathways for which the z-score is zero.

Ten statistically significant upstream regulators were identified by IPA® (Supplementary Table 5). The top activated upstream regulators were 3,5-dihydroxyphenylglycine and MLXIPL. The inhibited upstream regulators included FMR1 and LARP1. The top-ranked IPA Diseases and Disorders are shown in Figure 5 and the three top-ranked IPA Diseases and Disorders were *Immunological Disease*, *Organismal Injury and Abnormalities*, and *Inflammatory Response*.

**Figure 5:**
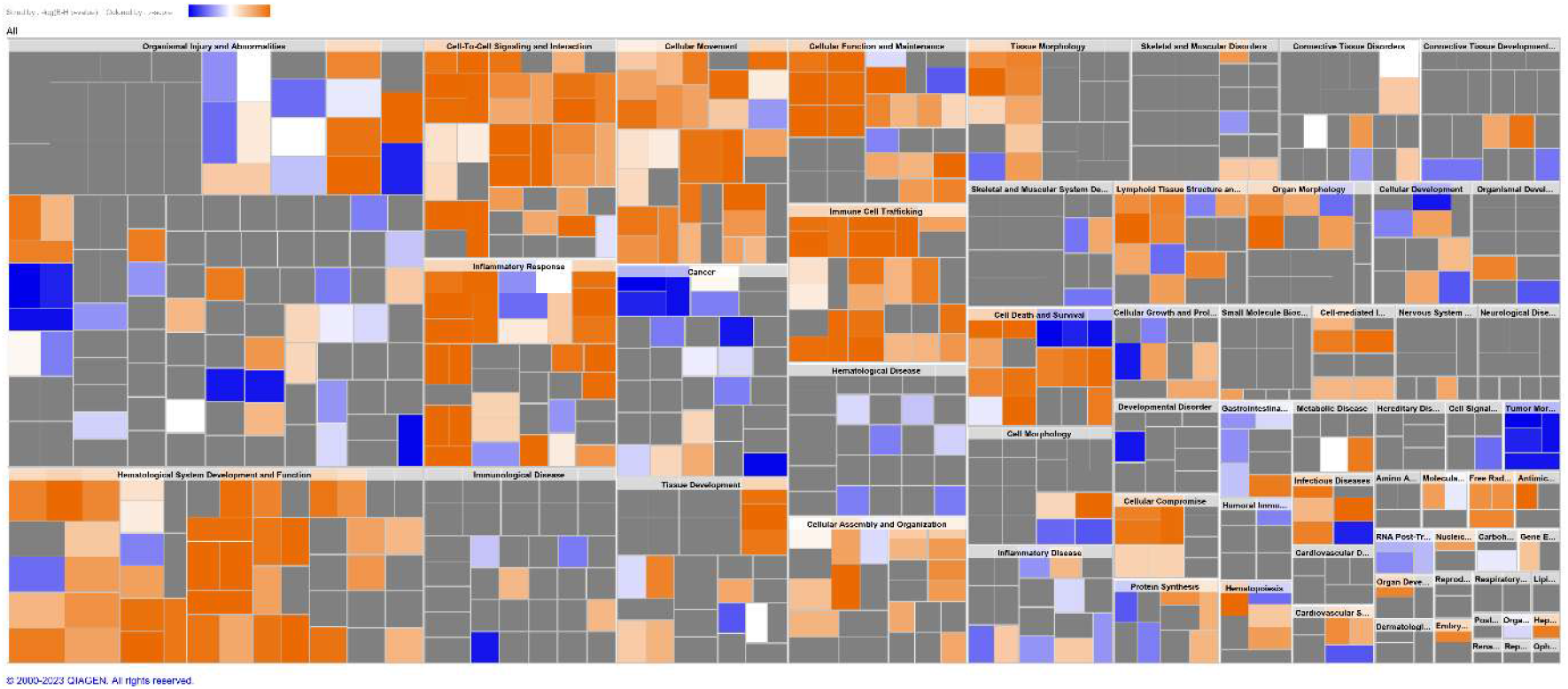
Heat map showing Diseases, Disorders and Biological functions identified using IPA®. The three most significant Diseases and Disorders were *Immunological Disease* (*P* value range: 0.04 – 0.0002), *Organismal Injury and Abnormality* (*P* value range: 0.04 – 0.0002), and *Inflammatory response* (*P* value range: 0.04 – 0.0002).

### Immunohistochemistry

Antibodies against common monocyte, dendritic cell and macrophage markers consisting of IBA-1, E-cadherin, CD163, CD301, CD90, and CD204, were applied to FFPE skin samples from ten cases of dogs with canine demodicosis and ten dogs with no history or histological evidence of skin disease (control group). All cases of canine demodicosis had significantly higher numbers of cells expressing positive immunolabelling for all antibodies (*P* ≤ 0.01), except for CD204 (P > 0.05), compared with the control group (Figures 6 and 7).

**Figure 6:**
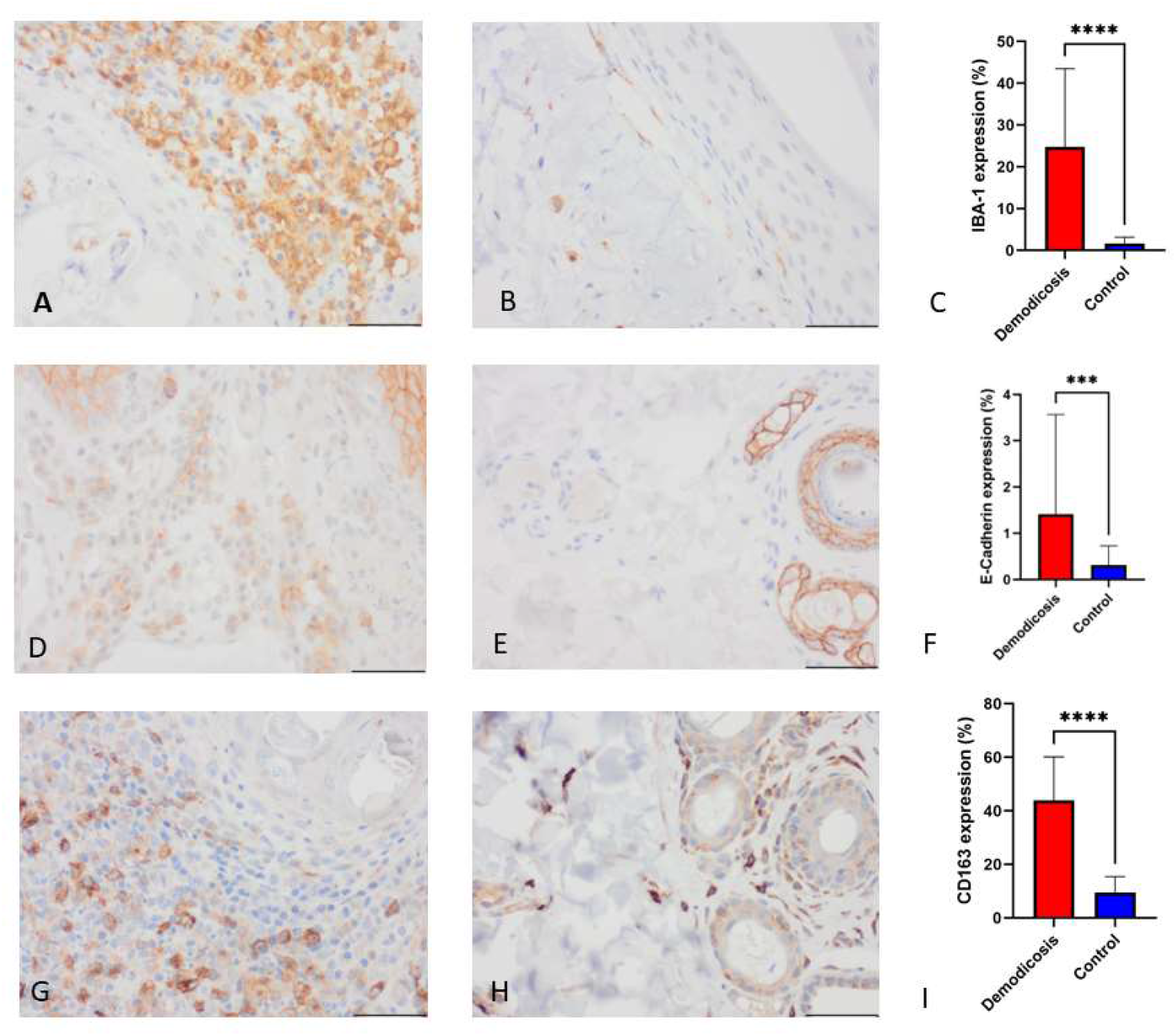
**A)** Skin from a canine demodicosis case, innumerable IBA1 positive immunolabelled cells are adjacent to a hair follicle containing a *Demodex* mite. **B)** Control skin, scattered IBA-1 positive immunolabelled cells within the dermis and extending into the outroot sheet of the hair follicle. **C)** Bar chart of percentage IBA-1 positively immunolabelled cells in cases of Demodicosis in comparison with controls (\*\*\*\**P* < 0.0001). **D)** Skin from a canine demodicosis case, moderate numbers of E-Cadherin positive immunolabelled cells are adjacent to two hair follicles that show normal E-Cadherin expression. **E)** Control skin, hair follicle epithelium is positively immunolabelled with E-Cadherin (normal expression). **F)** Bar chart of percentage E-Cadherin positively immunolabelled cells within the dermis in cases of Demodicosis in comparison with controls (\*\*\**P* < 0.001). **G)** Skin from a canine demodicosis case, numerous CD163 positive immunolabelled cells are adjacent to a hair follicle containing a *Demodex* mite. **H)** Control skin, scattered CD163 positive immunolabelled cells are present within the dermis. **I)** Bar chart of percentage CD163 positively immunolabelled cells in cases of Demodicosis in comparison with controls (\*\*\*\**P* < 0.0001).

**Figure 7:**
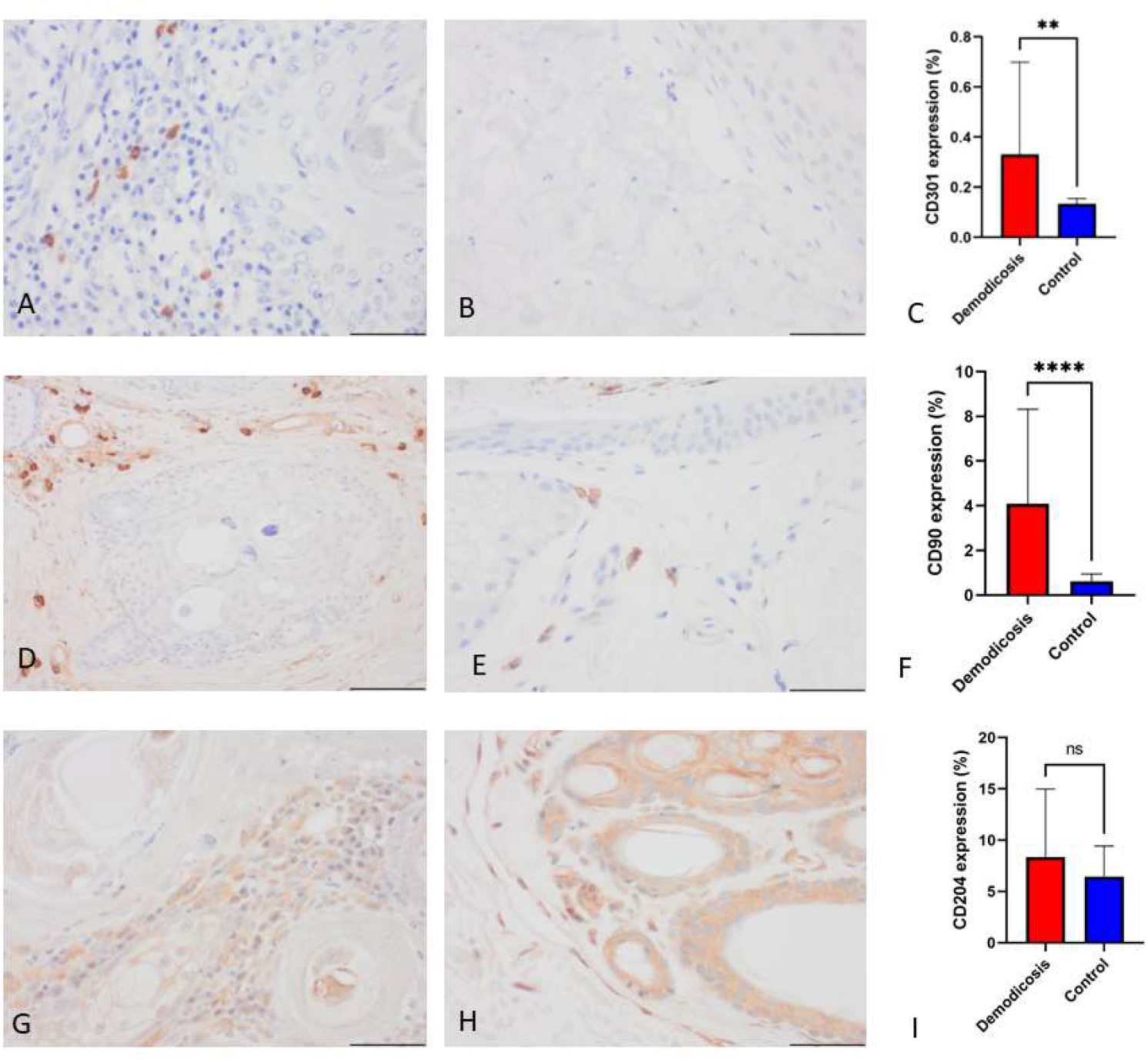
**A)** Skin from a canine demodicosis case, scattered CD301 positive immunolabelled cells are adjacent to a hair follicle containing a *Demodex* mite. **B)** Control skin, the dermis and hair follicle show no CD301 immunolabelled cells. **C)** Bar chart of percentage CD301 positively immunolabelled cells in cases of Demodicosis in comparison with controls (\*\**P* < 0.01). **D)** Skin from a canine demodicosis case, moderate numbers of CD90 positive immunolabelled cells surround a hair follicle that contains *Demodex* mites. **E)** Control skin, scattered CD90 positive immunolabelled cells are seen in the perifollicular areas. **F)** Bar chart of percentage CD90 positively immunolabelled cells within the dermis in cases of Demodicosis in comparison with controls (\*\*\**P* < 0.00001). **G)** Skin from a canine demodicosis case, numerous CD204 positive immunolabelled cells are adjacent to hair follicles. **H)** Control skin, scattered CD204 positive immunolabelled cells are present within the dermis surrounding hair follicles which also display non-specific cytoplasmic immunolabelling. **I)** Bar chart of percentage CD204 positively immunolabelled cells in cases of Demodicosis in comparison with controls (ns, *P* > 0.05).

## Discussion

The objective of this study was to assess the proteomic profiles of FFPE skin samples from dogs with canine demodicosis and compare these profiles with those from healthy canine controls. The aim was to provide further molecular insights into the pathogenesis of canine demodicosis. The study showed 154 protein groups to be significantly more highly abundant and 113 more lowly abundant in the demodicosis group compared to the control group. To further investigate these data, we employed two functional omics data analysis software tools, Metascape® and IPA®, which can be used to identify biologically relevant signalling and metabolic pathways, reconstruct molecular networks, predict the direction of downstream effects on biological and disease processes, and predict the activation and inhibition of upstream regulators(34, 35).

Metascape® analysis was used with both the high and low abundant proteins. Metascape® uses an overrepresentation analysis (ORA) method to assess the enrichment of pathways in a gene list compared to a background comparator list. It does not rank/prioritize proteins/genes based on statistical significance or differential expression (i.e., log_2_FC or *P*_adj._ value). Among the enriched ontology clusters in our study, we identified the *PI3 AKT pathway*. Although commonly associated with neoplasia, this pathway also plays a critical role in inflammation. It regulates Toll-like receptor (TLR) activity and NF-kB signalling in macrophages, leading to their polarization into M2 immunosuppressive macrophages.(39, 40). The presence of *Antigen Processing and Presentation of Exogenous Antigen Pathway* is not surprising given the presence of the exogenous *Demodex* mites. Similarly, the presence of *TNF Pathway*, *Leukocyte Activation* and *Leukocyte Cell-Cell Adhesion* is expected given the presence of inflammatory cells within the tissues assessed. The *TNF pathway* is essential for the immediate immune response and is involved the innate immune system through cell activation, proliferation, apoptosis and necrosis(41).

IPA® was used for gene set enrichment analysis (GSEA) in which proteins/genes can be ranked by statistical significance or expression difference (i.e., *P*_adj._ or log_2_FC), and statistically analysed for enrichment of specific pathways or other functional biological categories using detectable proteins/genes as the background comparator set. All gene symbol IDs for the significantly differentially abundant proteins were used for the IPA® analysis. This analysis identified four statistically enriched canonical pathways for canine demodicosis with biological processes and molecular functions similar to those identified by Metascape®. One of these pathways, the *Coronavirus Pathogenesis Pathway*, was incorporated into the Ingenuity® Knowledge Base in recent years(42) and has been observed in other studies of non-coronavirus infectious diseases(43). The three other significant enriched signalling pathways, *eIF2 Signaling, mTOR Signaling and Regulation of eIF4 and p70S6K Signaling*, are involved in regulation of initiation of transcription and translation in response to a range of stress stimuli both external and internal(44–46). These pathways have also been shown to be inextricably linked(46, 47) and all have been shown to be involved in the modulation of Toll-Like Receptors (TLR) (48, 49). Moreover, these pathways have been previously demonstrated to play a role in the pathogenesis of skin diseases such as rosacea, psoriasis, atopic dermatitis, and melanoma in both dogs and humans (50–54).

Cell stress, whether from internal or external sources, can cause improper protein folding resulting in the accumulation of misfolded or unfolded proteins in the endoplasmic reticulum (ER), known as ER stress. To combat this, the unfolded protein response (UPR) is initiated to restore normal ER function and reduce ER stress(52). The UPR reduces protein translation, increases degradation of misfolded/unfolded proteins, and increases the expression of ER chaperones and folding enzymes to improve protein folding. The eukaryotic translation initiation factor 2 (eIF2) signalling pathway is involved in the UPR, which regulates protein synthesis initiation. When eIF2 is phosphorylated, it inhibits eIF2B activity, leading to reduced protein synthesis and reduced protein folding load in ER-stressed cells. eIF2α selectively induces translation of activating transcription factor 4 (ATF4), which controls expression of adaptive genes that protect cells against ER stress. ER stress, through activation of ATF4 has been shown to affect proinflammatory cytokine production through modulating TLR signalling mostly notably through TLR2(48). ER stress via ATF4 also increases VEGF-A expression, stimulating angiogenesis and inflammatory lymphangiogenesis, which play a significant role in chronic skin inflammation(55). Both TLR2 and VEGFA have been shown previously to be upregulated in lesional skin from cases of canine demodicosis(13, 56). The findings suggest that ER stress-induced UPR signalling through the eIF2 pathway may contribute to the pathogenesis of canine demodicosis by upregulating TLR2 and activating the VEGFA-VEGFR2 signalling pathway.

There is growing evidence that the ER stress and UPR response is a modulator of immunity(57). Recent work has demonstrated that the ER stress and the UPR response are involved in regulating multiple immune cell types including T cells, B cells, DCs, macrophages, and myeloid derived suppressor cells (MDSCs) (57–60). More specifically ER stress and UPR responses have been shown to regulate differentiation, cytokine production, exhaustion, and apoptosis of CD8+ T cells as well as induce regulatory T cell plasticity. Additionally, the responses have been shown to support the survival of immunosuppressive MDSCs, trigger these cells to produce immunosuppressive iNOS, ROS, and Arg-1, promote IL-10 secretion and promote an immunosuppressive M2 macrophage phenotype(61–64).

In our previous gene expression study, we observed an increase in regulatory T cells and detected a gene expression pattern indicative of the presence of immunosuppressive MDSCs (myeloid-derived suppressor cells) and M2 macrophages in cases of canine demodicosis(56). Based on the pathways highlighted in both the Metascape® and IPA® analyses, specifically the PI3 AKT pathway and eIF2 signalling pathway, which are associated with the polarization of macrophages towards the M2 phenotype, we investigated the presence of M2 macrophages in the lesional skin of dogs with demodicosis. To accomplish this, we employed immunohistochemical markers commonly used for identifying dendritic cells and macrophages on the formalin-fixed paraffin-embedded (FFPE) tissue. In canine demodicosis, we observed a marked infiltration of cells positively immunolabelled with IBA-1, indicating the proliferation of Langerhans cells (LC), interstitial dendritic cells, and macrophages. Furthermore, we observed a significant infiltration of E-Cadherin positive (cytoplasmic and membranous) immunolabelled cells in the dermis, which confirm an increase in LCs and their mobilisation from the epidermis to the dermis. Research in the field of Leishmania, a protozoan parasite, has shown that LCs increase IL-10 and regulatory T cells (65). It is therefore possible that LCs are playing a role in promoting an immunosuppressive environment in canine demodicosis through similar pathways of IL10 production and induction of regulatory T cells. We also found an increase in CD90 positive cells in the demodicosis group, indicating proliferation of interstitial dendritic cells. Additionally, we observed a significant increase in CD163 and CD301 positive immunolabelled cells, which are common markers for M2 macrophages in the demodicosis group(66–69). M2 macrophages are known to produce immunosuppressive cytokines IL-10 and TGF-β together with growth factor VEGF(70). The immunohistochemical findings of significant infiltrations of CD163 and CD301 positively immunolabelled cells confirms the presence of M2 macrophages and supports their role in creating the immunosuppressive microenvironment seen in canine demodicosis.

The presence of M2 phenotype macrophages, along with the proteomic evidence of upregulated UPR response through *eIF2 signalling* pathway, supports the hypothesis that *Demodex* mites induce ER stress, resulting in the modulation of immunity towards an immunosuppressive, immune tolerance phenotype. Additionally, our previous findings of increased T regulatory cells and a gene expression profile supporting the presence of MDSC further reinforce this hypothesis, however further research into the canine MDSCs is required(56). Similar research into other pathogens, mostly protozoan infections, and mycobacterial diseases, support the ability of micro-organisms to induce ER stress, utilise the UPR response and polarise macrophages to the M2 immunosuppressive phenotype to avoid immune detection and allow for their proliferation(71, 72).

The protein known as mechanistic target of rapamycin (mTOR) plays a crucial role in the growth, proliferation, and differentiation of keratinocytes(73). mTOR is part of the PI3K/Akt signalling pathway, which, as noted earlier, plays a crucial role in regulation of inflammation. Previous studies investigating the role of mTOR signalling in helminth infections have shown that the pathway promotes M2 macrophage differentiation(69). This suggests the possibility of a role for mTOR signalling in regulating the M2 macrophages seen here in canine demodicosis. Disruption or dysregulation of mTOR, can also affect protein synthesis and thereby impact cell growth and proliferation leading to various skin diseases(74). Inflammatory cytokines such as IL-22, IL-17 and TNF can induce hyperactivation of the mTOR pathway, resulting in enhanced keratinocyte proliferation and reduced differentiation, as observed in the human skin disease psoriasis(50). A common histological change in canine demodicosis is epidermal and follicular epithelial hyperplasia, it could be postulated, given the current study, that this change is influenced by mTOR hyperactivation(4, 75). mTOR has also been implicated in the pathogenesis of rosacea, a human skin disease that is often compared with canine demodicosis due to increased populations of *Demodex* mites in lesional skin(76). Cathelicidin, LL-37, an antimicrobial peptide, which is increased in skin of rosacea patients, activates the mTOR pathway by binding TLR2 and results in increased expression of cathelicidin itself in keratinocytes. Also, it has been shown that cathelicidin again through mTOR signalling induces NF-kB activation resulting in cytokines and chemokines characteristic of rosacea and topical application of rapamycin, an mTOR inhibitor, results in improved clinical signs (76). Upregulation of TLR2 has also been shown in canine demodicosis, which may result in activation of the mTOR pathway thereby contributing to the cytokines and chemokines produced. mTOR has also been shown to be involved in angiogenesis in rosacea patients through increased cathelicidin and increased VEGF expression(77). While VEGFA expression has been shown to be increased in canine demodicosis its role in disease development and progression has not yet been elucidated(56).

The eIF4 and p70S6K signalling pathway is positively regulated by mTOR, and constitutes the main pathway for cell proliferation, survival, differentiation, and angiogenesis. Various stimuli, such as growth factors and cytokines, activate this pathway by triggering a phosphorylation cascade involving PI3K, Akt, PDK1 and mTOR(46). Activation of the PI3K/Akt/mTOR/p70S6K pathway impacts angiogenesis by the upregulation of VEGF(78). Dysregulation of this pathway was observed in the current study, however, its role and the role of VEGF in canine demodicosis remains to be clarified.

IPA® enrichment analysis identified immunological disease, organismal injury and abnormalities, and inflammatory response as the most significant underlying diseases and disorders. This is consistent with what would be expected for an inflammatory disease due to an ectoparasite.

This study has some limitations, such as the small number of significantly different protein groups (267) found between the disease and control groups. However, it is the first proteomic profiling study of skin samples from dogs with canine demodicosis. The skin samples used in this study were formalin-fixed paraffin-embedded (FFPE). Although formalin fixation causes significant cross-linking among proteins and other biomolecules, several studies have shown that there is a significant overlap in the number and identities of proteins between FFPE and frozen tissue(79–81). This indicates that there is an equivalence in proteomic profile obtained from fresh-frozen and FFPE tissue specimens by shotgun proteomics. Another limitation of this study is the utilization of distinct cohorts for the proteomic and immunohistochemical investigations, which was necessitated by the availability of only small tissue samples from clinical cases. The limited amount of available tissue is a persistent constraint in research endeavours. However, although this aspect can be perceived as a limitation, it also serves as a strength, as it not only demonstrates a proteomic profile indicative of M2 macrophages in ten cases but also confirms the presence of cells exhibiting M2 macrophage markers in an additional ten cases.

## Conclusion

The aim of this study was to gain molecular insights into the pathogenesis of canine demodicosis by employing proteomics and pathway enrichment analysis. Our findings revealed significant dysregulation in several interconnected pathways that regulate transcription and translation initiation in response to various internal and external stress stimuli. Furthermore, our pathway enrichment analysis using the proteomic profile identified immunological disease, organismal injury and abnormalities, and inflammatory response as the most significantly associated diseases and disorders. Immunohistochemistry confirmed the presence of M2 macrophages in lesional skin of canine demodicosis. Overall, our study suggests that *Demodex* mites trigger ER stress inducing an UPR response, leading to the proliferation of M2 macrophages, which, in turn, contributes to an immunosuppressive microenvironment, promoting the *Demodex* survival and proliferation.

## CRediT authorship contribution statement

Pamela A. Kelly, Rory Breathnach: Funding acquisition. Pamela A. Kelly, Rory Breathnach: Conceptualization. Pamela A. Kelly: Investigation, Writing − original draft. Pamela A. Kelly, Caitriona Scaife, Gillian P. McHugo, Susan Peters: Data curation. Pamela A. Kelly, Caitriona Scaife, Gillian P. McHugo: Formal analysis. Pamela A. Kelly, Caitriona Scaife, Gillian P. McHugo, David E. MacHugh, Susan Peters, Marion Stevenson: Methodology, Validation. Pamela A. Kelly, Gillian P. McHugo, Caitriona Scaife, Susan Peters, Marion Stevenson, Jennifer S. McKay, David McHugh, Irene Lara-Saez, Rory Breathnach: Writing − review & editing.

## Acknowledgement

The author(s) acknowledge the assistance of the Purdue University Histology Research Laboratory, a core facility of the NIH-funded Indiana Clinical and Translational Science Institute”, specifically laboratory manager Ms MacKenzie McIntosh, for the optimisation and staining of sections for E-Cadherin, CD163, CD90 and CD204. We also thank Dr Jamie Timmons (Augur Precision Medicine) and Dr Mark Ziemann (Deakin University) for valuable advice concerning the functional omics data analyses.

