## Supplementary figures 1-4 for "Unveiling the Role of Endoplasmic Reticulum Stress Pathways in Canine Demodicosis"

**Supplementary Figure 1:** *EIF2 Signaling Pathway* from Ingenuity® Pathway Analysis. Differentially expressed molecules in the pathway are highlighted with pink outlines. Increased or decreased expression is indicated by shades of orange and blue, respectively. Molecules with a double outline containing a gradient of these colours indicate groups or complexes of genes that may be significantly perturbed. A legend is available at https://qiagen.secure.force.com/KnowledgeBase/articles/Basic_Technical_Q_A/Legend.

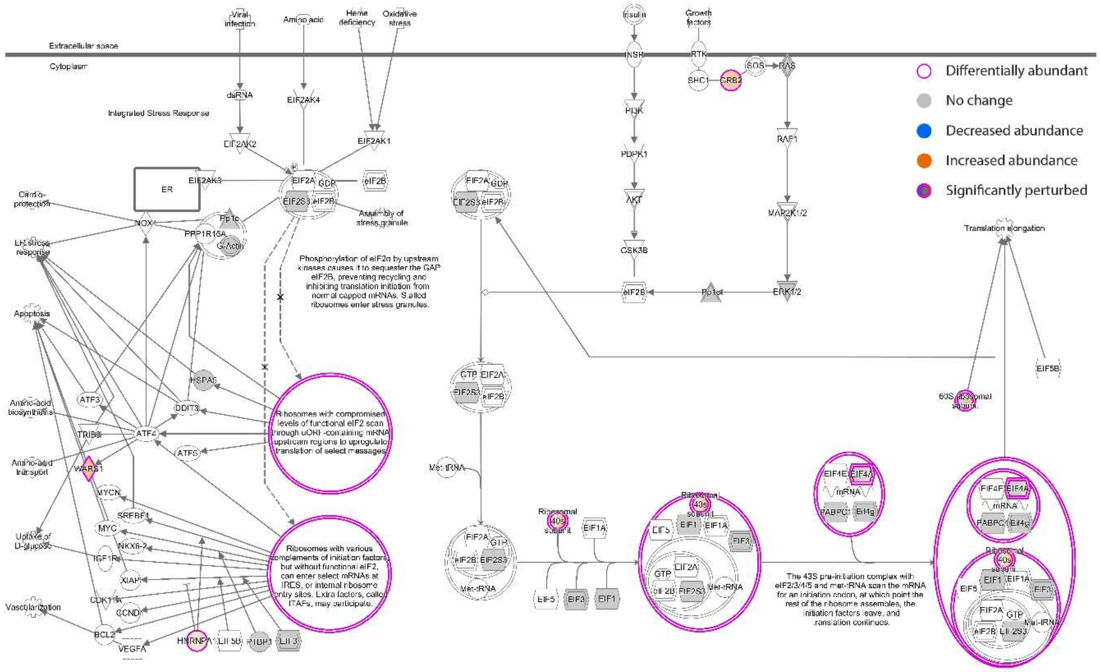


© 2000-2023 QIAGEN. All rights reserved.

**Supplementary Figure 2:** *Coronavirus Pathogenesis Pathway* from Ingenuity® Pathway Analysis. Differentially expressed molecules in the pathway are highlighted with pink outlines. Increased or decreased expression is indicated by shades of orange and blue, respectively. Molecules with a double outline containing a gradient of these colours indicate groups or complexes of genes that may be significantly perturbed. A legend is available at https://qiagen.secure.force.com/KnowledgeBase/articles/Basic_Technical_Q_A/Legend.

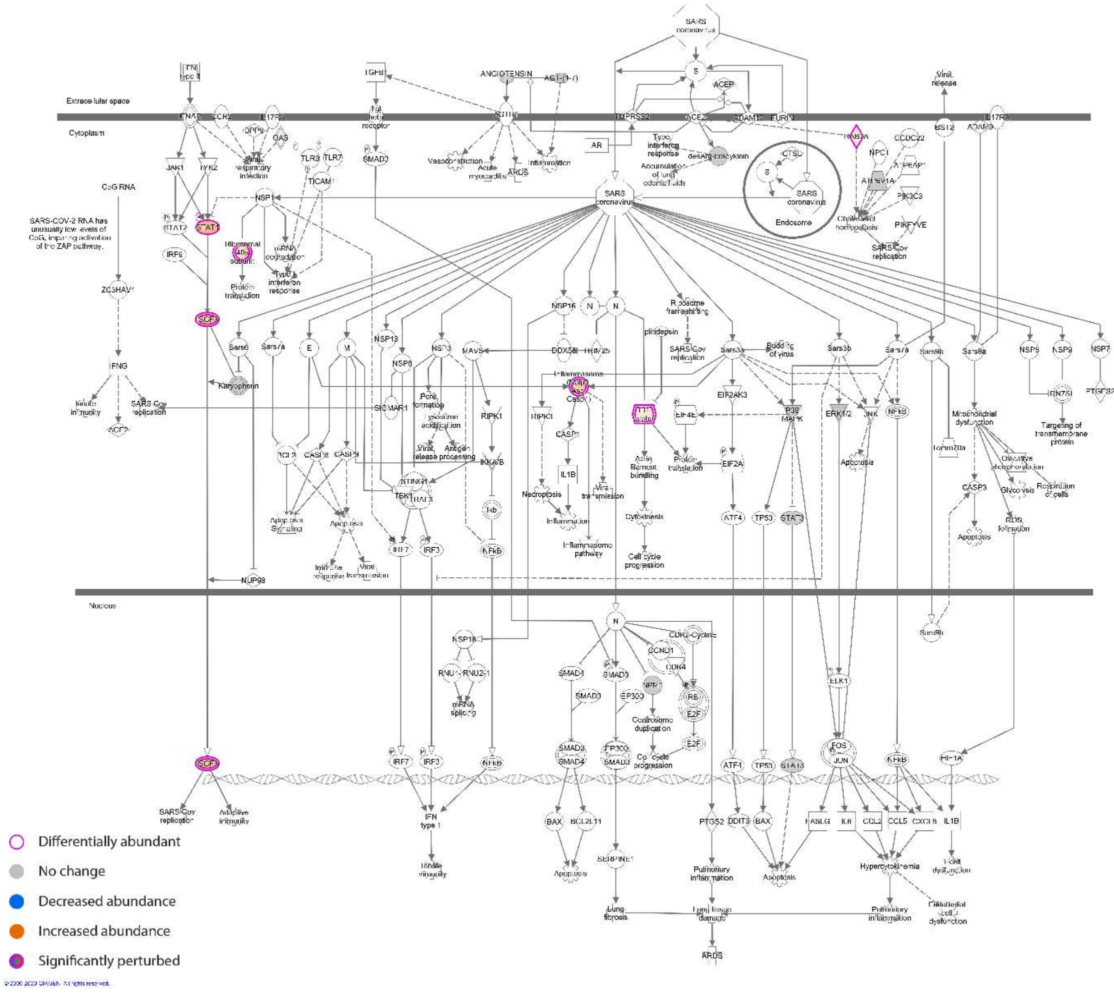

**Supplementary Figure 3:** *mTOR Signaling Pathway* from Ingenuity® Pathway Analysis. Differentially expressed molecules in the pathway are highlighted with pink outlines. Increased or decreased expression is indicated by shades of orange and blue, respectively. Molecules with a double outline containing a gradient of these colours indicate groups or complexes of genes that may be significantly perturbed. A legend is available at https://qiagen.secure.force.com/KnowledgeBase/articles/Basic_Technical_Q_A/Legend.

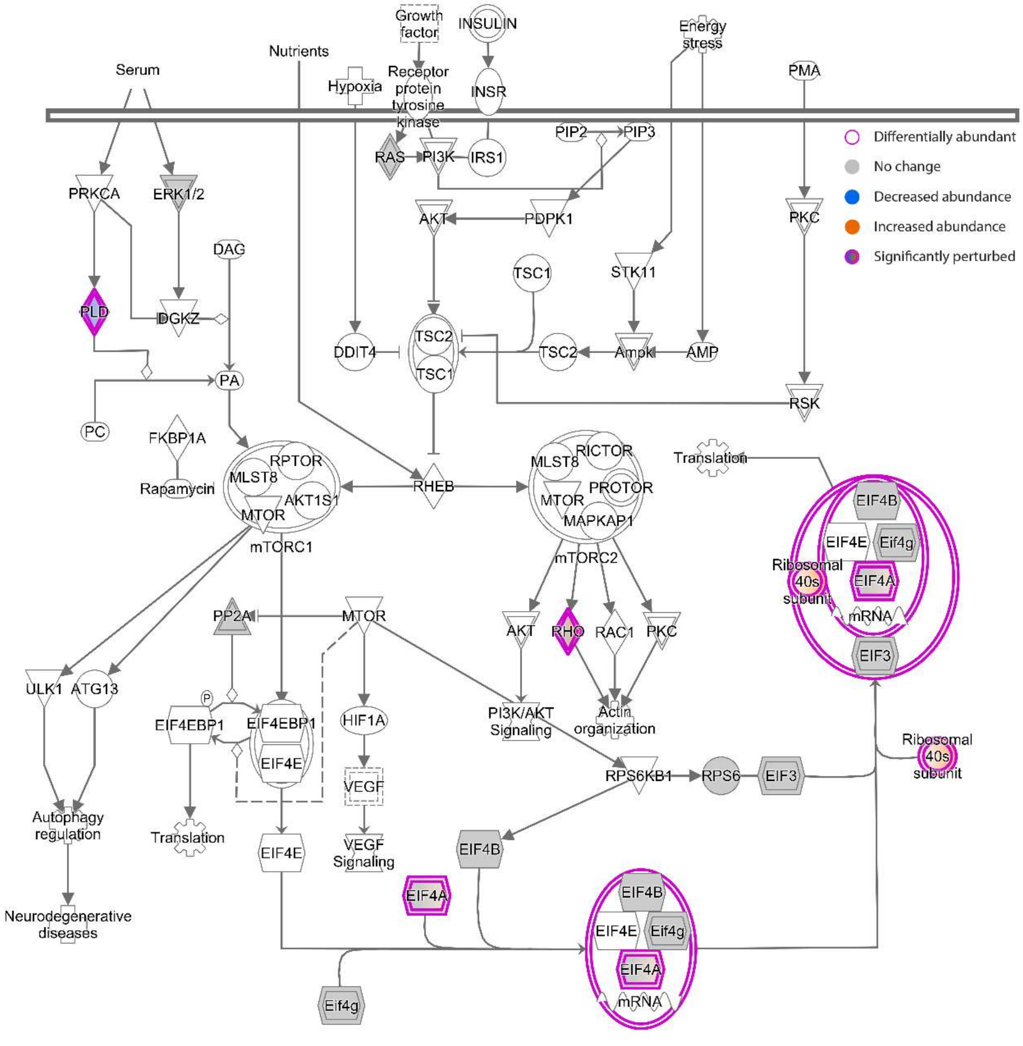


© 2000-2023 QIAGEN. All rights reserved.

**Supplementary Figure 4:** *Regulation of eIF4 and p70S6K Signaling Pathway* from Ingenuity® Pathway Analysis. Differentially expressed molecules in the pathway are highlighted with pink outlines. Increased or decreased expression is indicated by shades of orange and blue, respectively. Molecules with a double outline containing a gradient of these colours indicate groups or complexes of genes that may be significantly perturbed. A legend is available at https://qiagen.secure.force.com/KnowledgeBase/articles/Basic_Technical_Q_A/Legend.

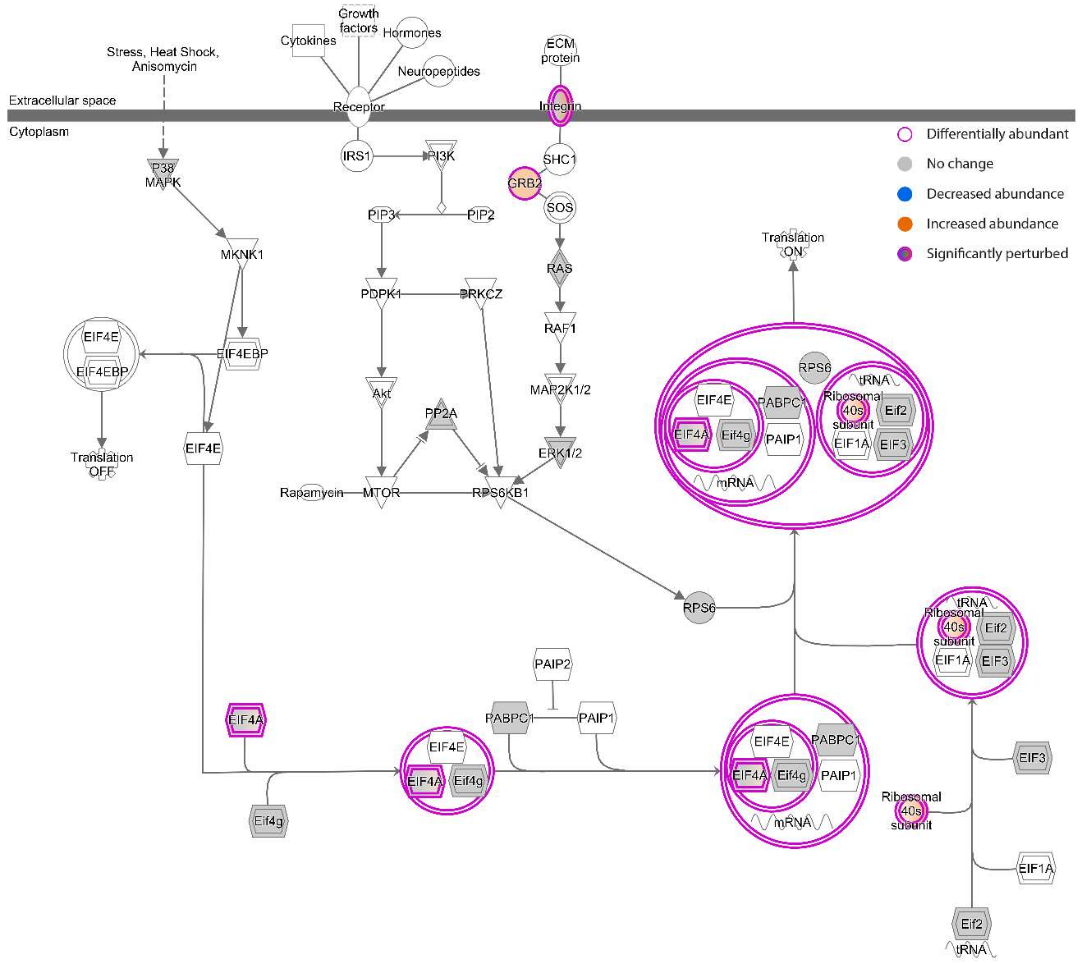


© 2000-2023 QIAGEN. All rights reserved.
