## Supplementary material 1 for "Unveiling the Role of Endoplasmic Reticulum Stress Pathways in Canine Demodicosis"

*Immunohistochemistry*

The antibodies applied are common cell markers used to identify different lineages of macrophage and dendritic cells in tissues. CD163 is considered a marker of M2 macrophages together with CD301 and CD204(36). IBA-1 is a dendritic cell marker that is expressed by Langerhans dendritic cells, interstitial dendritic cells, and macrophages. E-Cadherin is specific for Langerhans dendritic cells. CD90 is a marker of interstitial dendritic cells.

*IBA-1 and CD301*

Antigen retrieval was performed using heat-induced epitope retrieval (HIER), sections were treated at full pressure with an Access Retrieval Unit (Menarini, London, UK) in Sodium Citrate buffer (pH 6). The sections were then rinsed in Tris-buffered saline (TBS) buffer with Tween (pH 7.5).

The sections were treated for 5 min at RT with 3% hydrogen peroxide in PBS to quench endogenous peroxidase activity. After washing twice with TBS-Tween (pH 7.5), sections were incubated for 30 min at room temperature with the primary antibody CD301 at 1:50, IBA-1 at 1:1500, then washed with TBS-Tween (pH 7.5).

For primary antibody detection, the sections were incubated with EnVision+ System HRP Labelled Polymer Anti-Rabbit Secondary Antibody (Dako, Agilent, California, USA) for 30 min at RT. This was followed by a wash with TBS-Tween (pH 7.5) two 5-min incubations with 3,3'-diaminobenzidine (DAB) substrate-chromogen (EnVision+ System, Dako). Sections where then rinsed twice with $ddH_2O$ for 5 min. Tissues were counterstained using Gill's haematoxylin and mounted using DPX mounting media (Cellpath, Powys, UK) and coverslips for long-term storage.

*CD90, CD163, E-cadherin and CD204*

The paraffin embedded tissue blocks were sectioned at 4 µm thickness onto charged microscope slides and incubated at 57°C for 45 min. After incubation, microscope slides were deparaffinized and rehydrated through several changes of xylene and 100% ethanol, then rinsed in $ddH_2O$ before undergoing antigen retrieval. Heat-activated antigen retrieval was used for slides stained with E-cadherin (Cell Signaling Technology Inc., Massachusetts, USA) and CD163 (Novus Biologicals, Colorado, USA). Slides were placed in EDTA buffer in a decloaking machine heated to 95°C for 20 min. Once cooled to 60°C, tissues were rinsed with 1× Tris-EDTA (TE) buffer (pH 8) and placed on an intelliPATH automated slide stainer (Biocare Medical, California, USA). Enzymatic antigen retrieval was utilized for slides to be stained with CD90 (Novus Biologicals, cat. no: AF2067) and CD204 (Novus Biologicals, cat. no: NBP1-00092). Proteinase K was dispensed onto slides, which were incubated for 5 min, then rinsed with TE buffer.

Following retrievals, all slides were blocked with 3% $H_2O_2$ for 5 min. Tissues to be stained with CD90 and CD163 were rinsed and blocked with 2.5% Normal Horse serum (Vector Laboratories, California, USA) for 20 min. In addition, 2.5% Normal Goat serum (Vector Laboratories) was used for the same duration on tissues to be stained with E-cadherin and CD204. Primary antibodies were applied to the respective slides and incubated: CD90 for 60 min at 1:100, CD163 for 30 min at 1:250, CD204 for 30 min at 1:100, and E-cadherin for 30 min at 1:200. Slides were then rinses with TE buffer (pH 8) and incubated with HRP secondary antibodies for 30 min; goat anti-rabbit (Vector Laboratories) for CD204 and E-cadherin; horse anti-mouse (Vector Laboratories) for CD163; horse anti-goat (Vector Laboratories) for CD90. After an additional TE buffer rinse (pH 8), the Vector ImmPACT DAB chromogen (Vector Laboratories) was applied for 5 min. Lastly, slides were counterstained with Gills II Haematoxylin, dehydrated in ethanol, cleared in xylene, and cover slipped using a resinous mounting media.
